# Phosphorylated Pol II CTD supports histone hitchhiking during transcription elongation

**DOI:** 10.64898/2026.11.16.766941

**Authors:** Lulu Bi, Yikai Tong, Yifan Yang, Xinxin Wang, Yaqian Li, Xizi Chen, Yanhui Xu

**Affiliations:** Fudan University Shanghai Cancer Center, Institutes of Biomedical Sciences, New Cornerstone Science Laboratory, State Key Laboratory of Genetic Engineering, Department of Biochemistry and Biophysics, School of Life Sciences, and Shanghai Key Laboratory of Medical Epigenetics, Shanghai Medical College of Fudan University, Shanghai 200032, China; Sycamore Research Institute of Life Sciences, Shanghai 201203, China; The International Co-laboratory of Medical Epigenetics and Metabolism, Ministry of Science and Technology, China, Department of Systems Biology for Medicine, School of Basic Medical Sciences, Shanghai Medical College of Fudan University, Shanghai 200032, China

## Abstract

Transcription requires RNA polymerase II (Pol II) to disrupt nucleosomes while preserving displaced histones for reuse. How these histones remain available as the polymerase advances is incompletely understood. Here we combine optical trapping and fluorescence imaging to follow Pol II–nucleosome encounters in a simplified mammalian elongation system. Displaced histone particles frequently move with the advancing elongation complex, a behavior termed histone hitchhiking. Phosphorylation of the Pol II C-terminal domain (CTD) reduces nucleosomal blocking and supports histone retention during carriage. Structural predictions suggest that phosphorylated CTD repeats engage basic histone surfaces through variable, charge-complementary contacts. Carried particles can subsequently transfer to upstream DNA, and a broader combination of elongation factors increases upstream transfer. Complexes already carrying a histone particle show increased blocking at a second nucleosome. Other encounters leave one particle on upstream DNA while another moves with the complex, suggesting that transfer of one particle may enable capture and carriage of the next during successive nucleosome traversal. These observations support a model in which reversible contacts with the phosphorylated CTD help Pol II capture and carry histone particles as they dissociate from downstream DNA, keeping them available for upstream transfer and subsequent chaperone-assisted reassembly. Our findings extend the role of the Pol II CTD beyond factor recruitment to histone handling, linking nucleosome traversal to histone preservation.

## Introduction

Transcription by RNA polymerase II (Pol II) progresses from initiation through elongation to termination as regulatory factors dynamically associate with and dissociate from the polymerase [1]. The C-terminal domain (CTD) of its largest subunit, RPB1, provides a phosphorylation-regulated platform that coordinates these interactions throughout the transcription cycle [2]. TFIIH-associated CDK7 phosphorylates the CTD during initiation and promoter escape [1, 3, 4], whereas P-TEFb-associated CDK9 promotes pause release and productive elongation. CTD phosphorylation helps coordinate recruitment of co-transcriptional factors, including the 5′-capping machinery [5], splicing factors [6], elongation factors [7, 8], and histone-modifying enzymes such as SETD2 [9, 10]. Human Pol II has 52 CTD heptad repeats, of which approximately 17 support cell survival [11]; budding-yeast Pol II has 26 repeats, with eight sufficient for viability [12, 13]. Thus, roughly one-third of the repeats can sustain viability in these assays, but whether the extended array supports additional functions beyond factor recruitment remains unclear.

As the basic unit of chromatin, the nucleosome wraps approximately 147 base pairs (bp) of DNA around a histone octamer through extensive histone–DNA contacts [14]. These contacts fluctuate but collectively obstruct Pol II, which must gain access to the DNA template to continue transcription [15, 16]. Biochemical studies have shown that recovery of upstream histone–DNA contacts can be coupled to disruption of downstream contacts, preserving histones and restoring chromatin behind the advancing elongation complex (EC) [17, 18]. FACT [19–21], SPT5 [22], SPT6 [23], and IWS1/Spn1 [24, 25] provide histone-binding or chromatin-restoration activities that could contribute to this process. Structural studies have captured snapshots of intermediates in downstream DNA unwrapping, upstream DNA rewrapping and FACT-assisted nucleosome disassembly and reassembly [26–29]. These structures reveal local configurations for histone transfer around Pol II. However, how the EC progresses through chromatin while retaining displaced histones after they lose their original DNA contacts remains unclear.

## Results

### A simplified elongation system supports histone hitchhiking

To follow the fate of histones during nucleosome traversal, we established a simplified elongation system in which Pol II–nucleosome encounters could be monitored over time with fewer potentially redundant contributions from additional elongation factors. This provided a starting point for observing the behavior of the polymerase and displaced histones and then testing which components contribute to traversal and histone retention. We purified mammalian Pol II, human elongation factors, and histone chaperones, and combined optical trapping with multicolor fluorescence imaging (Fig. 1A and Fig. S1). DNA held between two optically trapped beads provided a template on which we followed elongation complexes through ATTO647N-labelled RNA 5′ ends (red) and histone particles through AF488-labelled H3.3 (blue). Force measurements and spatial mapping supported nucleosome assembly on the starting DNA template (Fig. S2A, B; Supplementary Note 1).

**Figure 1.**
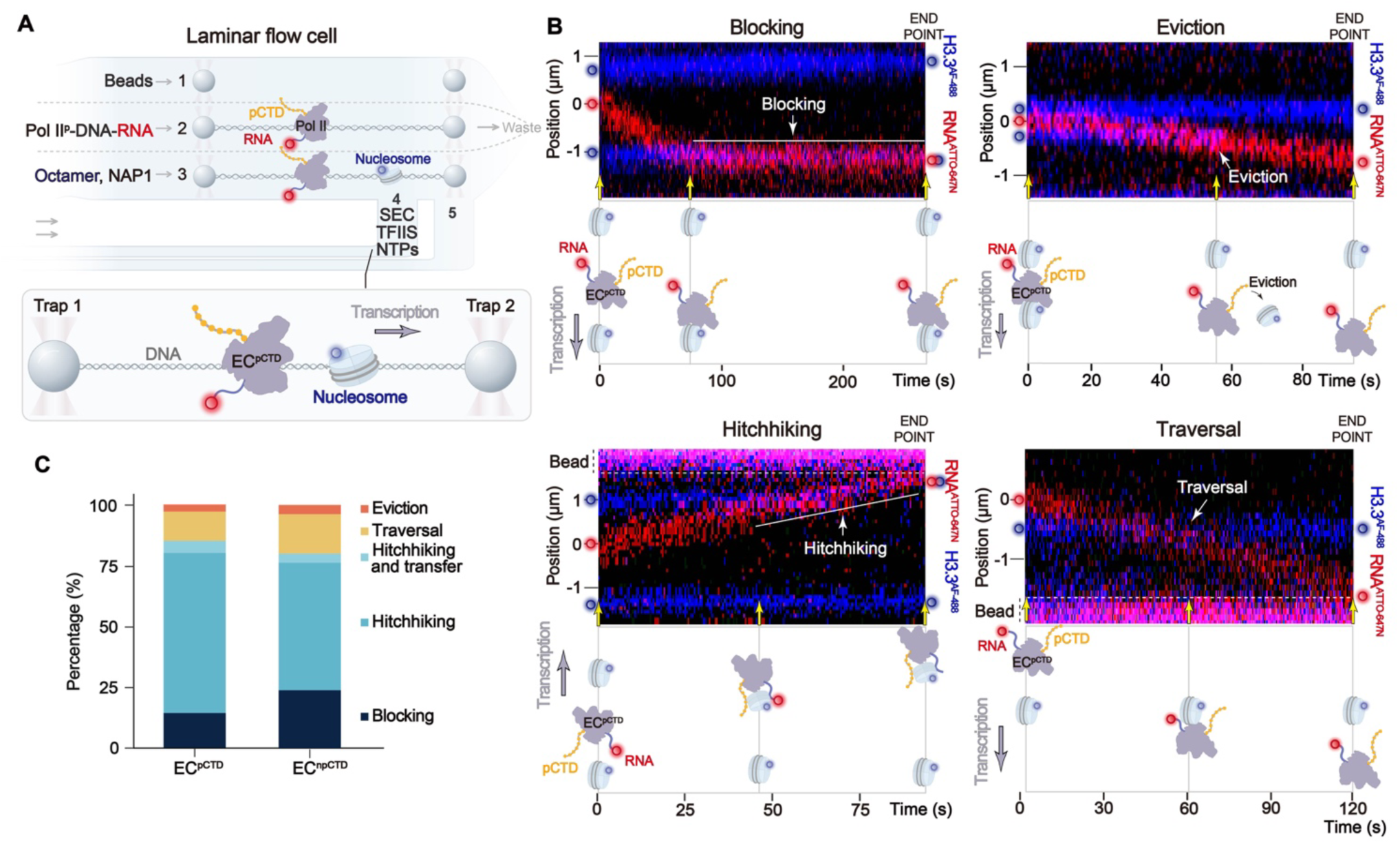
Single-molecule observation of histone hitchhiking during nucleosome traversal. **A,** Optical-trapping and fluorescence-imaging workflow for nucleosome assembly and transcription on a 16.2-kb DNA tether. EC^pCTD^ and EC^npCTD^ contain pre-phosphorylated and initially non-phosphorylated Pol II, respectively; both reactions contain TFIIS and the four-subunit SEC lacking P-TEFb. **B,** Representative EC^pCTD^ encounters showing blocking (upper left), eviction (upper right), hitchhiking (lower left) and traversal (lower right). ATTO647N-labelled RNA, red; AF488-labelled H3.3, blue. Red and blue circles indicate EC and histone-particle positions, respectively, at the beginning and end of each record. Schematics below depict the states at key time points indicated by yellow arrows. These display conventions apply throughout unless otherwise indicated. **C,** Encounter-outcome distributions for EC^pCTD^ (n = 141 records) and EC^npCTD^ (n = 80). Resolved carriage is divided into continued hitchhiking and hitchhiking followed by upstream transfer; event definitions are given in Methods.

We initially supplied Pol II with TFIIS, which promotes recovery from backtracking and can stimulate nucleosome traversal [30, 31]. On naked DNA, TFIIS alone supported slow elongation with prolonged pauses (Fig. S3A). The super elongation complex (SEC), which brings together P-TEFb and four additional subunits including AFF1/4, ELL1/2/3, ENL/AF9 and EAF1/2, associates with Pol II to promote elongation [32]. Addition of a four-subunit SEC containing EAF2, ELL2, ENL and AFF4 supported sustained movement along DNA, with a mean segmental elongation rate of approximately 2.3 kb min^−1^, comparable to rates supported by combinations of other elongation factors [33] and within the range reported in mammalian cells [34] (Fig. S3B, C). This four-subunit complex lacked P-TEFb and is referred to hereafter as SEC unless otherwise specified. It allowed elongation to be stimulated without adding P-TEFb kinase activity during the encounter. For the initial observations, Pol II CTD was pre-phosphorylated with P-TEFb and then transferred to the TFIIS–SEC elongation condition. We refer to these complexes as EC^pCTD^.

We first followed EC^pCTD^ as they approached individual nucleosomes and observed four types of behavior upon nucleosome encounter (Fig. 1B). In some records, the EC reached the nucleosome but failed to pass it within the observation window (blocking). In others, the histone-particle signal was lost (eviction). A third type showed the histone particle moving with the advancing EC (hitchhiking). In a subset of these trajectories, the histone particle subsequently stopped on DNA while the EC continued downstream, revealing histone transfer to upstream DNA after carriage (hitchhiking and transfer) (Fig. S3D, E). We also observed that some ECs appeared to pass a nucleosome that remained near its initial position on DNA, placing a histone particle upstream of the advancing EC (traversal), either after a pause (pause and traversal) or with no resolved pause (direct traversal) (Fig. 1B). Traversal without a resolved hitchhiking interval could reflect direct recapture by upstream DNA [28] or brief hitchhiking followed by upstream transfer below the assay’s spatial resolution of 0.1 µm. Here, histone particle denotes a fluorescently tracked H3.3-containing histone complex, which could be an H3–H4 tetramer, hexamer or octamer.

Among 141 EC^pCTD^ records, 20 resulted in blocking (14.2%), 4 in eviction (2.8%), 93 in hitchhiking (66.0%), 7 in hitchhiking and transfer (5.0%), and 17 in traversal (12.1%) (Fig. 1C). Hitchhiking (70.9% in total) was therefore the most frequent encounter behavior in this simplified system, suggesting that carriage could preserve displaced histones during polymerase passage and keep them available for upstream recovery. We therefore asked which component of the simplified EC sustains this association and allows carriage to bridge nucleosome traversal and histone recycling.

### CTD phosphorylation reduces blocking while carriage persists without phosphorylation

Although several elongation factors have established histone-binding activities, our simplified reaction contained only Pol II, TFIIS and the four-subunit SEC. TFIIS and these SEC subunits lack a well-characterized histone-binding module that would readily explain the observed carriage. We therefore considered whether electrostatic contacts could provide an alternative means of retention. As nucleosomal DNA unwraps, basic histone surfaces previously contacted by the negatively charged DNA backbone become exposed. A negatively charged component of the EC could engage these surfaces and retain the particle as it dissociates from downstream DNA. TFIIS and the SEC subunits contained relatively few extended acidic regions (Fig. S4). The CTD was a particularly attractive candidate because it remains covalently linked to Pol II as associated factors exchange. Its 52 heptad repeats in human Pol II can acquire numerous phosphate groups, producing a flexible, negatively charged chain. Such a chain could engage exposed basic histone surfaces before the particle dissociates from downstream DNA and help capture it as the original contacts are lost.

We next tested ECs containing non-phosphorylated Pol II, termed EC^npCTD^, under the same TFIIS–SEC conditions (Fig. S5A, B). Blocking occurred in 19 of 80 EC^npCTD^ records (23.8%), compared with 14.2% in EC^pCTD^ (Fig. 1C). The lower blocking fraction after phosphorylation was consistent with pCTD engaging basic histone surfaces exposed during unwrapping, thereby reducing rebinding and rewrapping of downstream DNA that would oppose Pol II advance. However, hitchhiking remained frequent without CTD phosphorylation (Fig. S5C), suggesting that other components could provide overlapping histone-binding activity. We therefore considered the nascent RNA. As the EC advances, the growing nascent RNA provides a strongly negatively charged phosphate backbone, which could bind exposed histone surfaces and partly substitute for pCTD in retaining displaced histones.

### RNA removal reveals a contribution of CTD phosphorylation to histone retention

Because removing RNA before an encounter largely prevented forward progression, we examined how hitchhiking was maintained after capture (Fig. S6 and Supplementary Note 2). We first allowed an EC^npCTD^ to encounter a nucleosome and verified that the histone signal moved with the complex. We then transferred the tether into an RNase A-containing channel and monitored the associated particle (Fig. 2A and Fig. S7). Following RNase A treatment, loss of the signal from the 5′-labelled RNA was followed shortly afterwards by loss of the histone signal (histone loss) (Fig. 2B). In some trajectories, the two signals disappeared nearly simultaneously (Fig. S7B). Thus, removal of nascent RNA was followed by loss of carried histones on different timescales. Notably, trajectories showed that ECs remained on DNA after RNase A treatment and could resume transcription upon return to the elongation channel, including after loss of the histone signal (Fig. 2C and Fig. S7C; Supplementary Note 2). These observations support histone dissociation as the explanation for histone-signal loss while the EC remains on DNA.

**Figure 2.**
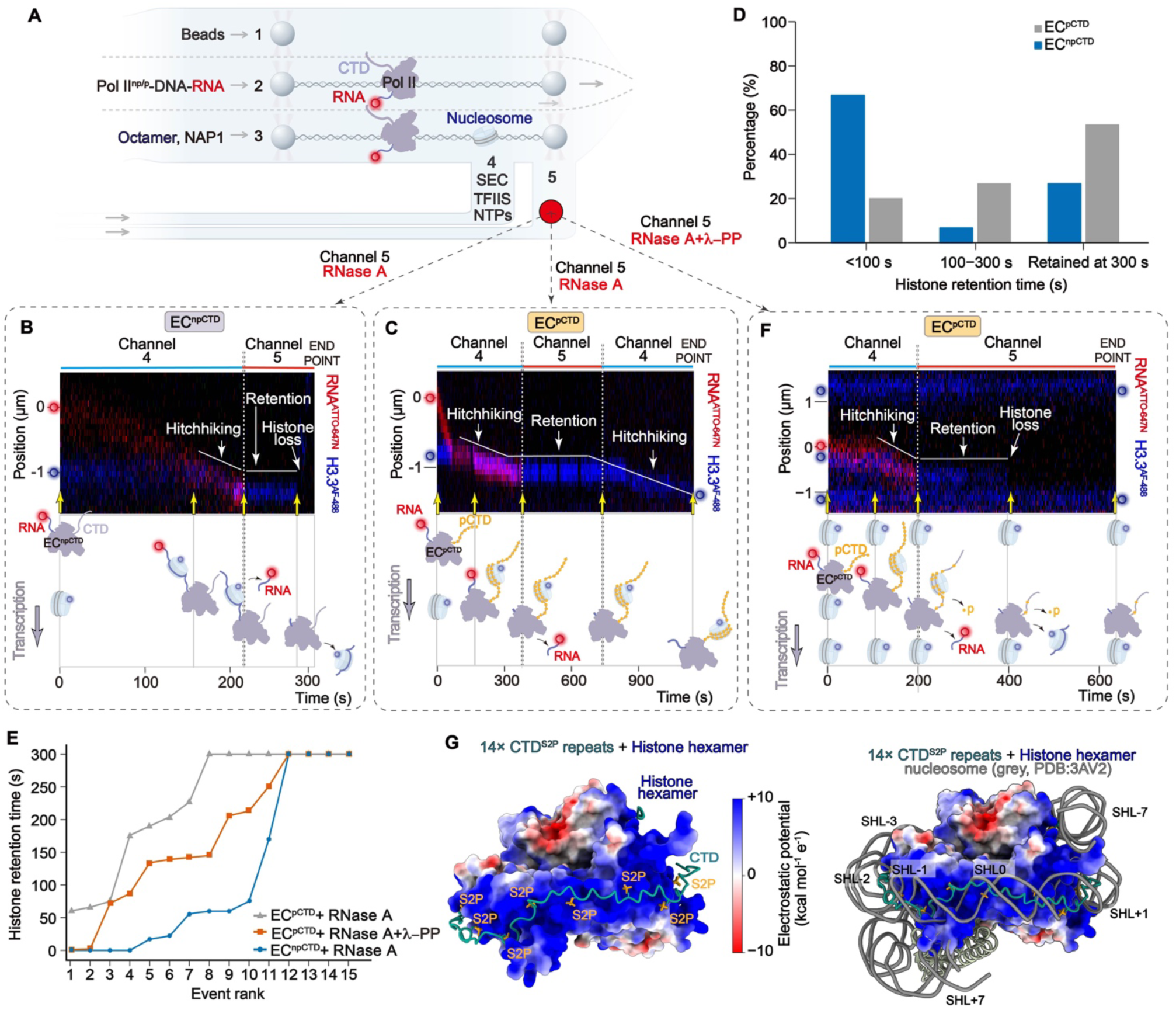
CTD phosphorylation supports histone retention after RNase A treatment. **A,** Workflow for RNase A or combined RNase A–λ-phosphatase (λ-PP) challenge of preformed EC–histone assemblies. **B,C,** Representative RNase A challenge records showing histone loss from EC^npCTD^ (B) and retention by EC^pCTD^, with carriage resuming on return to the elongation channel (C). **D,** Binned histone-retention times after RNase A treatment: <100 s, 100 ≤ t < 300 s, or retained at 300 s (n = 15 records per condition). **E,** Histone-retention times ranked independently for RNase A-treated EC^npCTD^ (blue circles), RNase A-treated EC^pCTD^ (grey triangles) and RNase A–λ-PP-treated EC^pCTD^ (orange squares; n = 15 each). Each point represents one record. Values of 300 s are right-censored; 0 s denotes loss within the time resolution. **F,** Representative trajectory showing loss of the carried histone particle following combined treatment with RNase A and λ-PP. **G,** Left, AlphaFold 3 model of a histone hexamer with 14 CTD repeats phosphorylated at Ser2. The histone surface is colored by electrostatic potential (red, negative; white, neutral; blue, positive; −10 to +10 kcal mol−1 e−1); CTD, teal; Ser2 phosphates, gold. Right, superposition with an H3.3 nucleosome (grey; PDB 3AV2). DNA was excluded from the prediction. SHL, superhelical location.

We then compared histone retention between EC^pCTD^ and EC^npCTD^ (Fig. 2D, E). In the EC^npCTD^ reaction, 10 of 15 particles (66.7%) were lost within 100 s and four (26.7%) remained detectable through the approximately 300-s observation window. In the EC^pCTD^ reaction, the corresponding counts were 3 (20.0%) and 8 (53.3%) of 15 particles. Residual retention in EC^npCTD^ could reflect RNA segments protected from cleavage by histone contacts or contributions from other EC components. EC^pCTD^ nevertheless retained histones more often throughout the observation window, supporting an additional contribution beyond accessible nascent RNA (Supplementary Note 2). In cells, RNA secondary structure formation and association with RNA-binding and processing factors may further restrict access to exposed histone surfaces [35, 36], and its cellular contribution remains to be determined.

### Phosphorylation-dependent contacts sustain histone carriage

To test whether phosphorylation-dependent contacts sustained histone retention after RNA digestion, we transferred preformed EC^pCTD^–histone complexes into channels containing RNase A together with λ-phosphatase (Fig. 2F). Combined treatment shortened histone retention relative to RNase A treatment of EC^pCTD^ alone, but histone loss remained slower than in RNase A-treated EC^npCTD^ (Fig. 2E, F and Fig. S7D). One possible explanation is that histone contacts limit access of λ-phosphatase to pCTD, delaying dephosphorylation and histone loss. Together with the EC^pCTD^ versus EC^npCTD^ comparison, these retention profiles support a role for phosphorylation-dependent contacts in sustaining histone carriage after removal of accessible RNA (Supplementary Note 2).

We next examined how a phosphorylated CTD chain might contact the exposed histone surface. Transcription can displace an H2A–H2B dimer and leave a hexasomal intermediate [28]. We therefore used AlphaFold 3 [37] to predict complexes of 7- or 14-repeat CTD segments phosphorylated at Ser2 with an H2A–H2B dimer, a histone hexamer lacking one H2A–H2B dimer, or a histone octamer (Fig. 2G and Fig. S8A–F). Across these combinations, CTD phosphate groups approached basic histone residues through variable contact configurations. This charge complementarity resembles that of nucleosomal DNA, whose principal histone-contact regions recur approximately every 10 bp at half-integer superhelical locations (SHL ±0.5 to ±6.5) [14]. In the representative superposition, 2-4 pCTD repeats approximately span a histone-surface region covered by 10 bp of nucleosomal DNA, suggesting less localized electrostatic contacts than those formed by the DNA backbone (Fig. 2G, Fig. S8 and Supplementary Note 3). In these predictions, Ser2 was phosphorylated in every repeat, but histone-contacting segments could be separated by segments that did not contact the surface. Together, the enzyme challenges and structural predictions suggest a model in which phosphorylation-dependent contacts help maintain histone carriage.

### Elongation factors promote histone transfer to upstream DNA

To examine how retained histones return to upstream DNA, we scored the endpoints of encounters in which the EC passed the nucleosome and the histone particle remained detectable (Fig. 3A, right). Upstream transfer occurred through three observed routes: direct traversal, hitchhiking and transfer, and pause and traversal. Continued hitchhiking comprised particles still moving with the EC at the observation endpoint. Adding FACT and NAP1 to EC^pCTD^ modestly increased upstream transfer (25.0% versus 20.5% without added chaperones). A combination of elongation factors containing PAF1C, RTF1, ELOF1, SPT6, pre-phosphorylated DSIF, TFIIS, P-TEFb, FACT and NAP1 (EC^All^) increased upstream transfer to 55.0%. With SEC additionally present (EC^All+SEC^), the corresponding fraction was 53.3%. Thus, a combination of elongation factors promoted transfer to upstream DNA, with similar observed fractions in the presence and absence of SEC.

**Figure 3.**
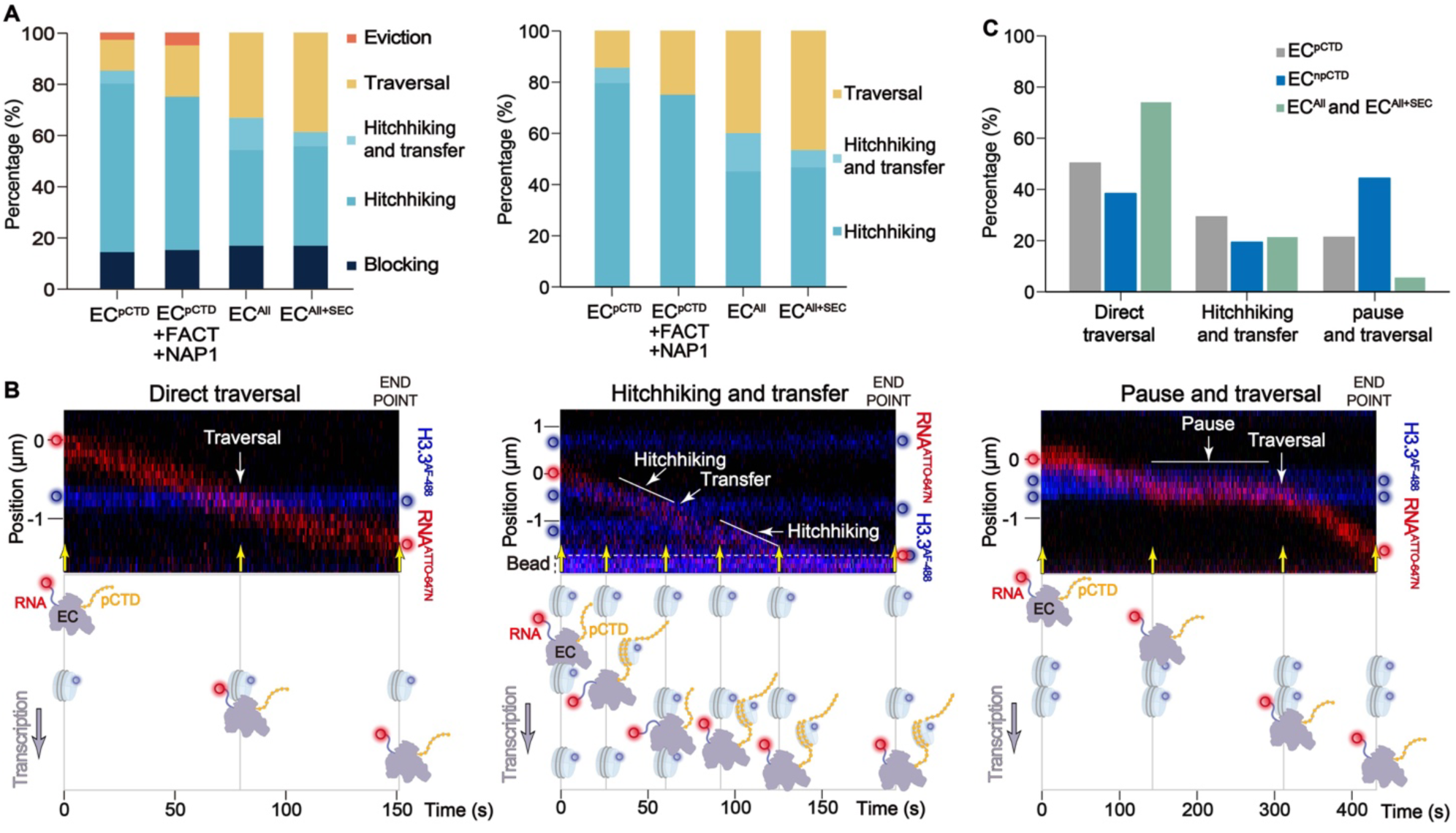
Elongation factors promote upstream transfer through distinct routes. **A,** Outcome distributions in EC^pCTD^, EC^pCTD^ plus FACT–NAP1, EC^All^ and EC^All+SEC^ reactions. Left, all recorded encounters (n = 141, 20, 24 and 18, respectively). Right, encounters remaining after exclusion of blocking and eviction (n = 117, 16, 20 and 15, respectively). Percentages are normalized separately for each condition in each panel. Upstream transfer comprises traversal and hitchhiking and transfer. All reactions used pre-phosphorylated Pol II; factor compositions are given in Methods. **B,** Representative kymographs and schematics of direct traversal (left), hitchhiking and transfer (middle), and pause and traversal (right). **C,** Route composition among upstream-transfer events in EC^pCTD^ (n = 24), EC^npCTD^ (n = 16) and pooled EC^All^ and EC^All+SEC^ reactions (n = 19).

We then compared the three routes among upstream-transfer events (Fig. 3B, C). The EC^pCTD^ reaction showed 50.0% direct traversal, 29.2% hitchhiking and transfer, and 20.8% pause and traversal. The corresponding proportions were 37.5%, 18.8% and 43.8% in the EC^npCTD^ reaction, and 73.7%, 21.1% and 5.3% in the pooled EC^All^ and EC^All+SEC^ reactions. Thus, transfer in the simplified EC^pCTD^ and all-factor conditions more often accompanied direct traversal or followed resolved hitchhiking, whereas EC^npCTD^ more often paused before traversal and upstream transfer. This comparison supports the role of pCTD in unwrapping nucleosomal DNA and suggests that phosphorylation and the all-factor mixture facilitate progression towards transfer. The shift towards direct traversal in the all-factor conditions is also compatible with factor-assisted histone transfer, rather than requiring unassisted recapture by upstream DNA alone.

Together, these results show that a combination of elongation factors increases the fraction of retained histone particles transferred to upstream DNA after nucleosome passage. Transfer occurred both after hitchhiking and without a detectable mobile intermediate, establishing hitchhiking followed by transfer as one route for histone preservation (Fig. 3B, C). Histone-binding modules and acidic or phosphorylated regions in these factors may assist both hitchhiking and upstream transfer (Fig. S4).

### Sequential nucleosome encounters reveal changes in particle carriage

We next asked how carrying one histone particle affected the next encounter. We followed EC^pCTD^ that captured a first particle and then encountered a second downstream nucleosome while still carrying the first. Among 11 trajectories, ECs stopped at the second nucleosome in 54.5% of records (Fig. 4A, D). Therefore, blocking was more frequent at the second encounter, when the EC already carried a histone particle, than at the first encounter (Figs. 1C and 4D). One possible explanation is that the first particle occupies part of the pCTD chain, leaving fewer contacts available to facilitate nucleosomal DNA unwrapping, engage the next nucleosome, and capture its displaced histones. CTD length and the distribution of phosphorylated repeats might therefore influence the capacity to accommodate successive particles (Supplementary Note 3).

**Figure 4.**
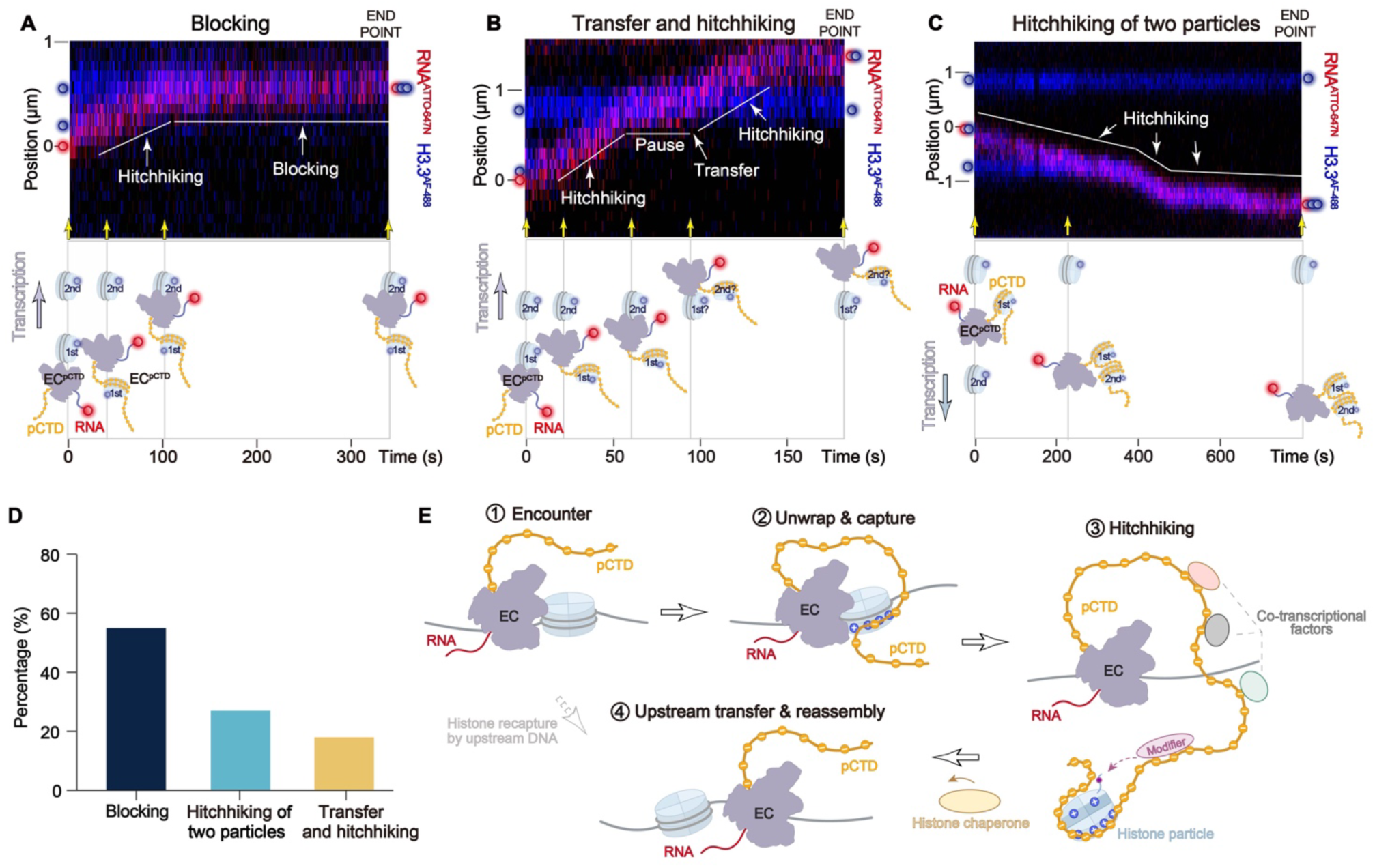
Sequential nucleosome encounters and a model for histone recycling. **A–C,** Representative trajectories of sequential nucleosome encounters by EC^pCTD^, showing blocking (A), transfer and hitchhiking (B), or hitchhiking of two particles (C) at the second encounter. Kymographs are shown above schematic interpretations; white lines highlight the trajectories. The identities of the histone particles after the encounter in B are unresolved. **D,** Outcome distribution for 11 second nucleosome encounters: blocking (n = 6), hitchhiking of two particles (n = 3) and transfer and hitchhiking (n = 2). **E,** Model of pCTD-assisted histone capture, hitchhiking and upstream transfer, with direct histone recapture by upstream DNA as a complementary route. Electrostatic contacts are proposed to support capture and carriage; histone modification and chaperone-assisted reassembly are shown as possible subsequent steps. Histone-particle composition remains unresolved. − and + denote negative charges on pCTD and positive charges on histone surfaces, respectively. EC, purple; pCTD, orange; DNA, grey; histones, light blue; RNA, red.

Two additional outcomes were observed (Fig. 4B–C). Firstly, one particle remained stationary on DNA at the second encounter site while another moved downstream with the EC, leaving the stationary particle upstream of the advancing enzyme (Fig. 4B, D). Although particle identities after the encounter were unresolved, a simple interpretation is that the EC transferred the first particle and then captured and carried the second (Supplementary Note 4). Secondly, ECs continued downstream while carrying both particles (Fig. 4C, D). These trajectories therefore revealed blocking, separation into one DNA-associated and one carried particle, and carriage of two particles as distinct responses to a second encounter.

### A model for pCTD-assisted histone recycling

Together, the single and sequential encounters support a model in which reversible pCTD contacts help Pol II capture and carry histone particles during transcription (Fig. 4E). As Pol II advances, nucleosomal DNA unwraps and exposes basic histone surfaces. By engaging these surfaces before the particle dissociates from downstream DNA, pCTD could partly compensate for lost histone–DNA contacts and reduce rewrapping that opposes Pol II advance (Supplementary Note 3). Continued engagement could allow pCTD to capture the displaced particle and carry it with Pol II until upstream DNA recapture becomes favorable. The particle can subsequently be transferred to upstream DNA, where histone chaperones might promote restoration of nucleosomal wrapping, as suggested by published work on FACT and upstream reassembly in our companion study [19, 21, 38] (Supplementary Note 5).

Transfer of the first particle could free pCTD contacts to engage the next nucleosome and capture and carry its displaced histones. Repeated exchange could allow the EC to transfer histones from successive downstream nucleosomes to upstream DNA, limiting the accumulation of displaced particles and, with subsequent reassembly, the extent of exposed DNA.

Histone-binding and acidic or phosphorylated regions in SPT6, IWS1 and SPT5 (Fig. S4) could make overlapping contributions to histone retention and transfer [22, 23, 25] (Supplementary Note 6). In the simplified reaction, pCTD-associated retention persists without these factors, suggesting that the covalently attached CTD could maintain histone retention as the associated factor complement changes. Engaging an extended histone surface could place an additional functional demand on CTD length beyond recruitment of cotranscriptional factors.

## Discussion

Structural studies have revealed downstream DNA unwrapping, upstream DNA rewrapping and FACT-associated intermediates that explain how histone contacts can be reorganized near Pol II [26–28]. Direct recapture requires upstream DNA to approach the exposed histone surface [28]. However, direct histone recapture by upstream DNA may be constrained by the bending cost of DNA looping [39] and by tension that opposes formation of a looped transfer intermediate [16] (Supplementary Note 7). Our trajectories reveal a complementary pathway in which displaced histone particles move with the EC before transfer to upstream DNA. Retention by a flexible pCTD could extend the time and range of positions available for DNA recapture. Together with upstream reassembly observed in the companion study [38], our findings support a model in which Pol II couples disruption of downstream nucleosomes to histone preservation and upstream chromatin restoration through a pCTD-supported mobile intermediate (Supplementary Note 5).

This model expands the role of CTD phosphorylation beyond recruitment of transcription-associated factors to the handling of histones during nucleosome passage. Retention near the EC may also bring these histones into proximity with CTD-associated chromatin modifiers. SETD2 provides a potential connection: its association with phosphorylated CTD and SPT6 contributes to co-transcriptional H3K36 methylation of upstream nucleosomes [9, 40, 41]. Such proximity raises the possibility of functional coupling between carriage and histone modification. This potential coupling links the proposed carrier function to the established regulatory platform of the CTD, suggesting how phosphorylation could coordinate nucleosome traversal with histone preservation, modification and chromatin restoration.

## Methods

### Protein expression and purification

RNA polymerase II (Pol II) was purified from *S. scrofa* thymus. Human TFIIS, P-TEFb, PAF1C, ELOF1, SPT6, DSIF and RTF1 were prepared as previously described [33]. Full-length constructs were used for all recombinant proteins, including the subunits of the multiprotein complexes.

The full-length open reading frames encoding the two human FACT subunits were cloned separately into modified pCAG vectors, with SUPT16H fused to an N-terminal protein A–SUMO tag. The two plasmids were cotransfected into Expi293F cells. After 72 h at 37 °C, cells were harvested and lysed for 30 min at 4 °C in lysis buffer containing 30 mM HEPES (pH 8.0), 300 mM NaCl, 0.25% CHAPS, 5 mM ATP, 5 mM MgCl_2_, 0.2 mM EDTA, 2 mM DTT, 10% (v/v) glycerol, 1 mM PMSF, 1 μg/mL aprotinin, 1 μg/mL pepstatin and 1 μg/mL leupeptin. The lysate was clarified by centrifugation at 15,000 rpm for 40 min at 4 °C, and the supernatant was incubated with IgG resin for 1.5 h. Bound proteins were released by on-column cleavage with ULP1 protease for 1 h at 4 °C. The eluate was further purified by anion-exchange chromatography on a Mono Q 5/50 GL column (Cytiva) equilibrated in buffer A (30 mM HEPES (pH 8.0), 30 mM NaCl, 2 mM DTT, 2 mM MgCl_2_ and 5% (v/v) glycerol). The proteins were eluted with a 0–100% gradient of buffer B (buffer A containing 1 M NaCl) over 30 column volumes. Peak fractions were pooled, aliquoted, flash-frozen in liquid nitrogen and stored at −80 °C.

The full-length open reading frame encoding human NAP1L1 (referred to as NAP1 throughout) was cloned into a modified pGEX-6P-1 vector to generate an N-terminal GST fusion containing an HRV 3C protease cleavage site. NAP1 was expressed in *E. coli* BL21(DE3) cells grown at 37 °C to an optical density at 600 nm (OD_600_) of approximately 0.6. Cultures were cooled to 16 °C, and protein expression was induced with 0.1 mM isopropyl β-D-1-thiogalactopyranoside (IPTG). After a further 16 h at 16 °C, cells were harvested and resuspended in lysis buffer containing 20 mM HEPES (pH 7.6), 500 mM NaCl, 0.05% CHAPS and 1 mM DTT. Cells were disrupted by high-pressure homogenization for 20 min at 4 °C, and the lysate was clarified by centrifugation at 15,000 rpm for 40 min at 4 °C. The supernatant was incubated with glutathione affinity resin for 1 h, and bound proteins were released by on-column cleavage with HRV 3C protease overnight at 4 °C. The eluate was further purified on a Mono Q 5/50 GL column (Cytiva) equilibrated in buffer A (20 mM HEPES (pH 7.6) and 1 mM DTT). The proteins were eluted with a 0–100% gradient of buffer B (buffer A containing 1 M NaCl) over 30 column volumes. Peak fractions were pooled, aliquoted, flash-frozen in liquid nitrogen and stored at −80 °C.

The full-length open reading frames encoding four human SEC subunits (AFF4, ELL2, ENL and EAF2) were cloned separately into modified pCAG vectors. EAF2 and AFF4 were fused to N-terminal protein A–SUMO–Spy and protein A–SUMO tags, respectively. The four plasmids were cotransfected into Expi293F cells. Cells were harvested 72 h after transfection and lysed for 30 min at 4 °C in lysis buffer containing 40 mM HEPES (pH 7.4), 300 mM NaCl, 0.25% CHAPS, 0.5 mM EDTA, 2 mM DTT, 10% (v/v) glycerol, 1 μg/mL aprotinin, 1 μg/mL pepstatin, 1 μg/mL leupeptin and 1 mM PMSF. The lysate was clarified by centrifugation at 15,000 rpm for 30 min at 4 °C, and the supernatant was incubated with IgG resin for 2 h at 4 °C. The resin was washed with wash buffer (30 mM HEPES (pH 7.4), 300 mM NaCl, 0.1% CHAPS, 2 mM DTT and 10% (v/v) glycerol) supplemented with 1 μg/mL aprotinin, 1 μg/mL pepstatin, 1 μg/mL leupeptin and 1 mM PMSF. Resin-bound proteins were dephosphorylated by overnight treatment with λ protein phosphatase (APExBIO, K1102) in wash buffer supplemented with 1 mM MnCl_2_ and lacking protease inhibitors. The resin was washed with inhibitor-free wash buffer, and the complex was released by on-column cleavage with ULP1 protease for 2 h at 4 °C. The eluate was diluted to a NaCl concentration of 200 mM and further purified by cation-exchange chromatography on a Mono S column equilibrated in buffer A (30 mM Bis-Tris (pH 6.4), 150 mM NaCl, 0.05% CHAPS, 2 mM DTT and 5% (v/v) glycerol). The complex was eluted with a linear NaCl gradient from 150 mM to 1 M over 30 column volumes. Fractions containing the purified SEC complex were pooled, concentrated using a concentrator with a 100-kDa molecular-weight cutoff, aliquoted and stored at −80 °C. For fluorescence labelling, SpyTag-bearing SEC was conjugated to Alexa Fluor 555-labelled SpyCatcher at a 1:1.5 molar ratio for 2 h at 4 °C. The estimated labelling efficiency exceeded 90%. The conjugated complexes were aliquoted, flash-frozen and stored at −80 °C.

### Preparation of nucleosomes

The coding sequences for human histones H2A, H2B, H3.3(K4C, C110A) and H4 were individually subcloned into pET-11b vectors, and the proteins were purified as previously described [42].

Lyophilized H3.3(K4C, C110A) was equilibrated at 22 °C in the dark for 1 h before reconstitution. A 1-mg sample was dissolved in freshly prepared labelling buffer (7 M guanidine hydrochloride (Gdn-HCl), 20 mM Tris-HCl (pH 7.0) and 5 mM EDTA) to a final protein concentration of approximately 100 μM. Disulfide bonds were reduced by adding tris(2-carboxyethyl)phosphine (TCEP; 0.5 M stock) to a final concentration of 1.25 mM, followed by incubation on a tube rotator at 22 °C for 2 h. Alexa Fluor 488 C5-maleimide (Thermo Fisher Scientific, A10254; 10 mM stock in DMSO) was added at a sevenfold molar excess relative to histone, with the final DMSO concentration kept below 10% (v/v). The reaction mixture was incubated in the dark at 4 °C for 16 h and then at 22 °C for 1.5 h. Insoluble aggregates were removed by high-speed centrifugation, and unbound dye was removed by extensive dialysis against buffer containing 7 M Gdn-HCl, 20 mM Tris-HCl (pH 7.5) and 10 mM DTT. Labelling efficiency was assessed by 18% SDS–PAGE.

Fluorescent histone octamers were reconstituted as previously described [42, 43]. Briefly, labelled H3.3 was mixed with unfolded wild-type histones H4, H2A and H2B at a molar ratio of 1:1:1.2:1.2. The mixture was dialysed at 4 °C against at least three changes of refolding buffer (2 M NaCl, 10 mM Tris-HCl pH 7.5, 1 mM Na-EDTA and 5 mM 2-mercaptoethanol). The assembled octamers were purified by size-exclusion chromatography on a Superdex 200 Increase 10/300 GL column (Cytiva) equilibrated in refolding buffer. Peak fractions containing histone octamers were pooled and stored at −80 °C.

### In vitro assembly of Pol II–DNA–RNA ternary complexes

To assemble the Pol II–DNA–RNA ternary complex, 8.1-kb DNA handles were prepared by digesting the corresponding plasmids with BsaI (New England Biolabs, R3733S). One cohesive end was filled in and biotinylated using Klenow fragment (3′→5′ exo−; New England Biolabs, M0212S) in the presence of biotin-11-dUTP (Jena Bioscience, NU-803). Fluorophore-labelled RNA and 5′-phosphorylated DNA oligonucleotides were purchased from Shanghai Generay Biotech Co., Ltd. The RNA–DNA hybrid (RDH) was prepared by mixing the 5′-phosphorylated template DNA strand (93Tem, 5′-CAACCGTTTGTGTTTTGCAAAGGTCTCGATATCTTTCTGGCTTGGGTTGGCTTTTCGCCGT GTCGTACATCATACTACTCCTACCAGGAAGCA-3′) with ATTO647N-labelled RNA (RNA24-647N, 5′-UUUUUUUUUUUUAUACGACACGGC-3′) at a DNA:RNA molar ratio of 1.5:1. The final RDH concentration was 5 μM, calculated from the RNA concentration. The mixture was incubated at 85 °C for 5 min, cooled to 25 °C at 1 °C per 60 s and held at 25 °C for 10 min.

The annealed RDH was incubated with Pol II at an RDH:Pol II molar ratio of 1:2.0 in a 20-μL reaction containing 4 μL of 0.5 μM RDH, 2 μL of Pol II, 0.5 μL of RNase inhibitor and 1 μL of 0.1 M DTT in 1× assembly buffer (20 mM HEPES (pH 7.5), 100 mM KCl, 5 mM MgCl_2_ and 10% (v/v) glycerol). The reaction was incubated at 25 °C with shaking at 350 rpm for 20 min. The 5′-phosphorylated non-template DNA strand (93Nts, 5′-ACCATGCTTCCTGGTAGGAGTAGTATGATGTACGACACGGCGAAAAGCCAACCCAAGCC AGAAAGATATCGAGACCTTTGCAAAACACAAACG-3′) was then annealed to the RDH-Pol II complex by adding 1.8 μL of 1 μM 93Nts. For reactions requiring Pol II phosphorylation, 0.8 μL of 10 μM P-TEFb and 0.5 μL of 100 mM ATP were added. The mixture was incubated at 30 °C with shaking at 350 rpm for 20–25 min, yielding an assembled Pol II–DNA–RNA ternary complex at a theoretical concentration of approximately 80 nM. Complexes for the EC^npCTD^ condition were prepared by omitting the Pol II pre-phosphorylation step.

For single-molecule experiments, the 8.1-kb biotinylated DNA handles were ligated to the upstream and downstream cohesive ends of the DNA scaffold in the ternary complex. Ligation was performed in a 20-μL reaction containing the assembled ternary complex, equimolar amounts of the DNA handles, 0.5 μL of T4 DNA ligase and an additional 0.5 μL of RNase inhibitor. After incubation at 16 °C for 1–2 h, the ligation mixture was kept on ice until use.

### Single-molecule optical tweezers and fluorescence imaging

Single-molecule manipulation and multicolour fluorescence imaging were performed using a C-Trap G2 optical tweezers system (LUMICKS) equipped with a five-channel microfluidic flow cell and a confocal microscopy module. All assays were performed at 28 °C. Trap separation was controlled by steering one trap with a piezoelectric mirror. Force was measured by back-focal-plane interferometry of the condenser lens using a position-sensitive detector. A computer-controlled stage enabled rapid, automated translation of the optical traps between channels of the laminar-flow cell for buffer exchange [44].

Before sample introduction, protein-containing channels were passivated with 1 mg/mL bovine serum albumin (BSA) in assay buffer for 10 min without flow. Unbound BSA was removed by flushing the channels with working buffer. Streptavidin-coated polystyrene beads (2.14 μm; Spherotech, SVP-20-5) were captured in dual optical traps in channel 1. The traps were moved to channel 2, which contained preassembled Pol II–DNA–RNA ternary complexes with biotinylated DNA handles at both ends, to form a tether between the beads. Single tethers were verified from their force–extension curves [45]. The tethered complex was then moved to channel 3, containing 1 nM Alexa Fluor 488-labelled histone octamers and 2 nM NAP1 in assembly buffer (20 mM HEPES (pH 7.5), 100 mM KCl, 5 mM MgCl_2_ and 10% (v/v) glycerol). After multiple fluorescent spots along the DNA tether were detected by real-time fluorescence scanning, the complex was transferred to channel 4 for transcription elongation assays. To investigate the mechanical properties of nucleosomes, we performed stretching experiments using optical tweezers at a constant trap-separation velocity of 0.2 μm s−1. Nucleosomal DNA unwrapping was identified from force transitions in the force–extension curves.

For the simplified condition, channel 4 contained 0.5 μM TFIIS and either 60 nM unlabelled SEC or 20 nM fluorophore-labelled SEC. In the EC^pCTD^ plus FACT–NAP1 reaction, 100 nM each of FACT and NAP1 were added to the simplified condition. DSIF was pre-phosphorylated separately before being added to the channel 4. For EC^All^, channel 4 contained 100 nM PAF1C, 120 nM SPT6, 400 nM RTF1, 1 μM TFIIS, 200 nM pre-phosphorylated DSIF, 1 μM ELOF1, 250 nM FACT and 50 nM NAP1. EC^All+SEC^ contained the same factors with 60 nM SEC added. Neither mixture contained IWS1. 50 nM P-TEFb was present in channel 4 in both broader-factor conditions, in addition to the treatment during complex preparation described above. Unless otherwise stated, transcription elongation assays were performed in elongation buffer containing 20 mM HEPES (pH 7.5), 100 mM KCl, 5 mM MgCl_2_, 10% (v/v) glycerol, 0.1 mg/mL BSA, 1 mM of each NTP (ATP, UTP, CTP, and GTP) and 0.1 U/μL murine RNase inhibitor. RNase A (New England Biolabs, T3018L) treatment was performed at a final concentration of 20 µg/mL in a reaction buffer containing 20 mM HEPES (pH 7.5), 100 mM KCl, 5 mM MgCl₂, 10% (v/v) glycerol, and 0.1 mg/mL BSA, omitting RNase inhibitors. For the co-treatment with λ-phosphatase (New England Biolabs, P0753L), the reaction was carried out in the supplier-provided buffer (50 mM HEPES, 100 mM NaCl, 2 mM DTT, 0.01% Brij 35, and 1 mM MnCl₂) with λ-phosphatase at a final concentration of 20 U/µL. During RNase A and combined RNase A–λ-phosphatase challenges, the flow cell was flushed with 0.1-bar pressure pulses lasting approximately 10 s at intervals of 1–2 min to remove released histones and reduce nonspecific histone–DNA reassociation.

Confocal line scans (kymographs) were acquired in channels 3, 4 or 5 by repeatedly scanning a laser line along the longitudinal axis of the DNA tether, using a pixel size of 100 nm, a dwell time of 0.1 ms per pixel and a total line time of 500 ms. Excitation wavelengths of 488, 561 and 638 nm were used to image Alexa Fluor 488-labelled histones, Alexa Fluor 555-labelled SEC and ATTO647N-labelled RNA, respectively. To avoid artifacts in extension calculations, the optical traps were held at a fixed position to maintain the DNA tether under a baseline tension of approximately 1 pN.

### Single-molecule data analysis

Force–extension measurements and binding kinetics were recorded using Bluelake software (LUMICKS). Kymographs and corresponding force–time traces were analysed using Lakeview software (LUMICKS). Trajectories of Alexa Fluor 488-labelled histones, Alexa Fluor 555-labelled SEC and ATTO647N-labelled RNA were automatically tracked using the built-in algorithms to determine their positions along the DNA tether over time. The elongation rate was calculated in accordance with the methodology described in a previous study [33]. To convert transcription-associated displacement into base pairs, the force–extension relationship of double-stranded DNA was measured by stretching the Pol II–DNA–RNA tether construct in the same reaction buffer and fitting the data to an extensible worm-like chain model. By fitting the force–extension curves obtained from 12 independent, singly tethered DNA molecules within this experimental setup, the apparent extension per base pair at a baseline tension of 1 pN was determined to be 0.278 nm/bp. Elongation rates and nucleosomal DNA-unwrapping forces are reported as the mean ± standard error of the mean (s.e.m.) for the indicated number of events. For the tracking of histone-binding dynamics during RNase A and λ-phosphatase treatments, histone retention was scored for up to 300 s after entry into the challenge channel. Particles still detectable at 300 s were classified as retained at this observation endpoint.

### Event classification and observation endpoints

Encounters were described as blocking, eviction, hitchhiking or traversal. Blocking denoted failure to pass the nucleosome within the observation window; eviction denoted histone-particle loss of unassigned cause. Hitchhiking denoted resolved co-movement of a histone particle with the EC. In the distributions, continued hitchhiking was labelled “Hitchhiking”, whereas resolved carriage followed by upstream transfer was labelled “Hitchhiking and transfer”. These two categories together represent encounters with resolved histone carriage. Traversal denoted passage without a resolved carriage interval, leaving the histone particle on upstream DNA. Traversal included direct traversal with no resolved pause and pause and traversal, in which passage followed a pause. Slow but resolved EC–histone co-movement followed by upstream transfer was classified as hitchhiking and transfer (Fig. S3D). Histone particle refers to the fluorescently tracked H3.3-containing histone complex; possible compositions include an H3–H4 tetramer, hexamer or octamer. H3.3 fluorescence does not determine stoichiometry or establish that nucleosomal DNA wrapping persists during carriage. For the comparison in Fig. S5C, blocking and eviction were excluded; the denominator was the sum of continued hitchhiking, hitchhiking and transfer, and traversal records. The pooled dataset included initially colocalized EC–histone records that subsequently co-moved (Supplementary Note 1). For the endpoint analysis in Fig. 3A, right, both blocked encounters and encounters with histone eviction were excluded; Fig. 3A, left, includes all recorded encounters. Upstream transfer included particles remaining on upstream DNA after traversal without resolved carriage or after hitchhiking; continued hitchhiking denoted carriage persisting to the observation endpoint. The denominator was the sum of continued hitchhiking, hitchhiking and transfer, and traversal records. Transfer-route composition in Fig. 3C used only upstream-transfer events, classified as direct traversal, hitchhiking and transfer, or pause and traversal. EC^All^ and EC^All+SEC^ records were pooled for this route comparison, but remained separate for the endpoint comparison in Fig. 3A. Histone signals retained through the approximately 300-s enzyme-challenge observation window were right-censored. Values recorded as 0 s denoted loss within the experimental time resolution. Ranked-duration and violin plots describe the observed records and do not provide uncensored dissociation lifetimes.

### AlphaFold 3 prediction of histone–CTD complexes

Complexes of histones and the Pol II C-terminal domain (CTD) were predicted using AlphaFold 3 [37]. Each input contained one CTD chain comprising seven or fourteen tandem YSPTSPS repeats (49 or 98 residues, respectively), with Ser2 in every repeat specified as phosphoserine. The CTD was modelled with an H2A–H2B dimer, an H2A–H2B–(H3.3–H4)₂ hexamer or an (H2A–H2B)₂–(H3.3–H4)₂ octamer. Histone constructs were derived from the resolved residues of the corresponding chains in the human H3.3 nucleosome structure [46] (PDB 3AV2), with unresolved terminal regions omitted. Each histone was supplied as a separate protein chain, and DNA was excluded.

Predictions were performed locally using AlphaFold 3 (v.3.0.0), except for the seven-repeat CTD–octamer complex, which was predicted using AlphaFold Server. Local runs used paired and unpaired multiple sequence alignments and structural templates, with five random seeds (1–5) and five models per seed, yielding 25 models per condition. For the seven-repeat CTD–octamer complex, five AlphaFold Server submissions used seeds 1–5 with structural templates enabled for all protein chains. Each submission generated five models, yielding 25 models in total.

Models without an AlphaFold clash flag were ranked by the CTD chain-specific interface predicted template modelling (ipTM) score. Ties were resolved sequentially by higher mean CTD–histone chain-pair ipTM, lower mean bidirectional CTD–histone predicted aligned error (PAE), higher overall ranking score and higher mean CTD atomic predicted local distance difference test (pLDDT) score. A complementary screen considered the consistency of CTD–histone contact patterns across seeds, contact statistics and interface-confidence metrics. Candidates from both screens were inspected manually to select representative conformations. Histone surface electrostatic potentials were calculated and visualized using UCSF ChimeraX [47] (Fig. 2G and Fig. S8).

## Acknowledgments

We thank the core facility at Fudan University for their support with data collection. We thank Drs. Shixin Liu and Yukun Wang at Rockefeller University for their comments on this study. Funding: This work was supported by grants from the National Key R&D Program of China (2021YFA1300100, 2022YFA1303200), the National Natural Science Foundation of China (32571496, 32271242, 32030055, 31830107, 31821002), the Basic Research Special Zone Plan of Shanghai (22TQ020), the XPLORER Prize (Y. X), the New Cornerstone Investigator Program (Y. X), and the National Program for support of Top-Notch Young Professionals (X.C.).

## Author contributions

L. B. and Y. T. performed the experiments and conducted data analysis with the help of Y. Y.; X. W. and Y. L. prepared the protein samples; X. C. and Y. X. designed the experiments, analyzed the data and wrote the manuscript; Y. X. supervised the project. Competing interests: The authors declare no competing interests.

## Data availability

All data are available in the main text or supplementary materials.

## Supplementary Information

### Supplementary Note 1. Characterization of the starting nucleosome substrate

Before monitoring transcription, we compared naked-DNA and nucleosome-loaded tethers, both containing preassembled Pol II–DNA–RNA ternary complexes. The loaded tethers exhibited force-induced DNA-release transitions near 20 pN, consistent with nucleosomal DNA unwrapping (Fig. S2A) [48]. Spatial mapping showed histone particles distributed along DNA in both phosphorylation conditions, whereas preloaded Pol II complexes were concentrated near the tether midpoint (Fig. S2B). The relative frequency of histone particles appeared slightly higher near the preloaded complex in the pCTD condition, raising the possibility that pCTD promotes local histone association during substrate preparation.

Some histone particles were initially colocalized with the preloaded complex and subsequently moved with the EC. The force and spatial measurements support nucleosome assembly on the starting substrate and are consistent with some initially colocalized particles being nucleosomes assembled near Pol II, below the approximate 0.1-µm spatial discrimination limit. Direct capture of histones by pCTD during preparation could also contribute.

### Supplementary Note 2. RNase and phosphatase treatments

RNase A treatment before a nucleosome encounter caused loss of the RNA signal and intermittent SEC association without sustained forward movement (Fig. S6). Nascent RNA can influence Pol II pausing and backtracking [49], providing a possible explanation for the reduced progression. We therefore applied the enzyme challenges after histone carriage had formed to examine how the captured particle was retained by the EC.

In an EC^pCTD^ trajectory, the RNA signal disappeared after RNase A treatment while the histone particle remained detectable (Fig. 2C). Returning the tether to the elongation channel led to resumed movement of the particle, showing that it remained associated with a complex capable of continuing transcription. In a separate experiment with fluorescent SEC, the carried histone signal disappeared after approximately 200 s of RNase A exposure, whereas the SEC signal persisted and resumed movement after return to the elongation channel (Fig. S7C). These trajectories show that an EC can remain on DNA after loss of the nascent-RNA signal and subsequently resume transcription. This is consistent with earlier single-molecule measurements of yeast Pol II elongation in the presence of RNase A [49].

The different retention profiles are consistent with overlapping contributions from RNA and pCTD (Fig. 2D, E and Fig. S7D). pCTD may engage exposed histone surfaces during DNA unwrapping and maintain these contacts after histone displacement. RNA could provide alternative or additional contacts. After RNA digestion, some particles could remain associated through pre-existing pCTD contacts or could be transferred from RNA to pCTD during the challenge.

Combined RNase A and λ-phosphatase treatment challenges an assembly formed with a phosphorylated CTD, whereas EC^npCTD^ captures histones without prior CTD phosphorylation. For particles captured by EC^pCTD^, subsequent phosphatase treatment does not imply complete CTD dephosphorylation. The intermediate retention profile after combined treatment (Fig. 2E) could reflect differences in how RNA- and pCTD-mediated contacts are disrupted. Cleavage of an accessible RNA segment can interrupt an RNA-mediated tether, whereas destabilizing a multivalent pCTD–histone association may require removal of the majority of contributing phosphates. Histone binding could also reduce phosphate accessibility, potentially delaying particle release.

### Supplementary Note 3. pCTD–histone contacts and the capacity for histone retention

Nucleosomal DNA contacts the histone core at 14 principal regions centered on half-integer superhelical locations, SHL ±0.5 to ±6.5 [14]. The CTD models suggest a related interaction principle in which multiple phosphate groups approach basic histone surfaces (Fig. 2G and Fig. S8). Histone-contacting stretches are separated by segments that do not contact the surface. Because every repeat in the model inputs was phosphorylated at Ser2, this separation reflects the chain conformation. Superposition with a nucleosome places these predicted contacts in the context of surfaces normally engaged by DNA.

These predicted pCTD–histone contacts could partly compensate for the loss of histone–DNA interactions, stabilizing a displaced particle and keeping it available for recapture by upstream DNA (Fig. 4E). Subsequent binding to DNA could replace the transient pCTD–histone contacts with histone–DNA interactions.

The repeat architecture raises the possibility that CTD length and the distribution of available contacts influence the capacity to retain one or more histone particles. A related geometric question is how many phosphorylated repeats are needed to contact a substantial portion of the exposed surface of an H2A–H2B dimer or histone hexamer. Human and budding-yeast Pol II contain 52 and 26 CTD repeats, respectively [2]. A 17-repeat human CTD supported cell survival in complementation assays [11]. Cell survival does not establish the repeat requirement for histone carriage.

### Supplementary Note 4. Molecular interpretations of sequential nucleosome encounters

The trajectories in Fig. 4B, in which one histone particle remains on DNA while another moves downstream with the EC, are consistent with two interpretations. In the exchange model, the first carried particle is transferred to DNA and the second is captured by the EC. Transfer could free histone-binding contacts for capture of the second particle. Alternatively, the first particle could remain associated with the EC as Pol II traverses the second nucleosome, whose histones remain on or return locally to DNA. This alternative requires the EC to handle the second nucleosome while retaining the contacts that carry the first particle.

### Supplementary Note 5. Relationship to the companion study

Both the present and companion studies observe displaced histone particles remaining associated with an advancing EC before returning to upstream DNA. The companion study establishes upstream nucleosome reassembly with mammalian ECs in a broader elongation-factor system, whereas our simplified system examines a candidate role for pCTD in histone retention [38]. The reassembly observed in the companion study, together with published work on FACT-mediated nucleosome reorganization, supports chaperone-assisted nucleosome reassembly as a possible continuation of histone transfer [19, 21].

The companion study uses H3.1-containing particles labelled on H4, whereas our assay tracks labelled H3.3. These differences, together with differences in DNA-template design, should be considered when comparing event frequencies.

### Supplementary Note 6. Potential contributions of other elongation factors

Other elongation factors may provide complementary contacts through acidic or phosphorylated regions (Fig. S4). Candidates include phosphorylated SPT5 and the acidic N-terminal region of SPT6. The latter binds histones, including the DNA-binding surface of H2A–H2B, and has a functional connection with FACT supported by genetic studies [23]. Beyond the factors tested here, IWS1 is another candidate, as its yeast homologue Spn1 binds H3–H4 through its acidic N-terminal region and interacts with Spt6 [24, 25]. Such interactions could help retain or transfer particles within the broader elongation-factor system. Our mixture comparisons do not isolate the contribution of SPT6 or any other individual factor. Defined factor combinations would help separate these contributions from direct CTD effects and changes in factor recruitment.

### Supplementary Note 7. DNA bending and tension in upstream histone recapture

DNA bending and tension may constrain the ability of upstream DNA to recapture displaced histones. In a reconstituted Pol II transcription system, applied tension reduced histone transfer to upstream DNA, consistent with inhibition of a looped intermediate [16, 39]. The transfer distances inferred in these experiments depend on DNA-loop geometry and flexibility [39]. Transcription-generated torsion may further influence nucleosome dynamics under rotational constraint, with its effects depending on polymerase rotation and the capacity of chromatin to accommodate torsional stress [50, 51]. These physical constraints motivate the possibility that retention by a flexible pCTD keeps a displaced particle available as Pol II advances, extending the time and range of positions over which upstream DNA can recapture it.

**Figure S1.**
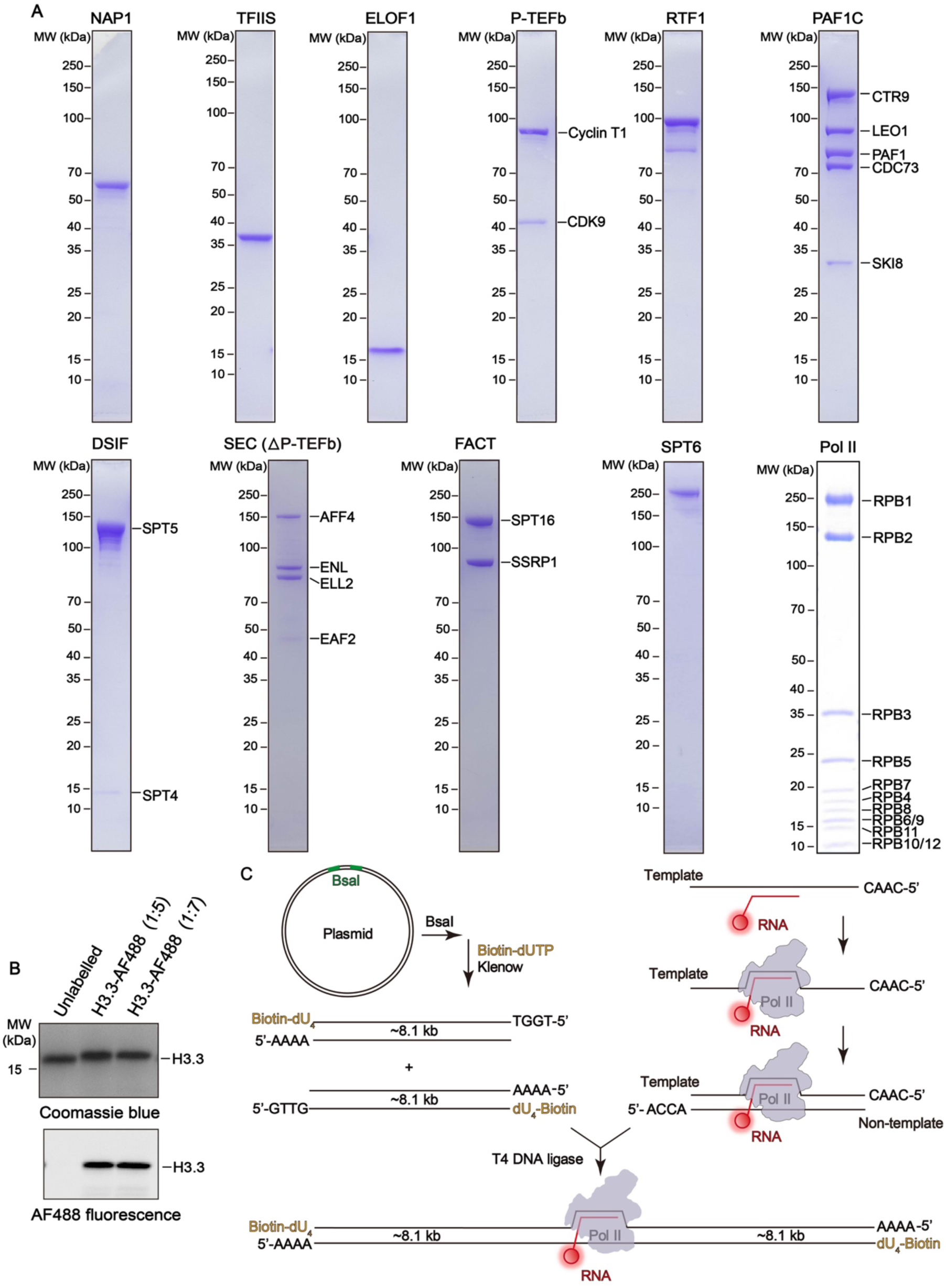
Protein purification, histone labelling and assembly of Pol II–DNA–RNA tethers. **A,** Coomassie blue-stained SDS–PAGE gels of the indicated purified proteins and complexes. SEC contains AFF4, ENL, ELL2 and EAF2. **B,** Coomassie blue staining (top) and AF488 fluorescence (bottom) of unlabelled and labelled H3.3 at the indicated H3.3:AF488 molar ratios. **C,** Assembly of a Pol II–DNA–RNA ternary complex between two approximately 8.1-kb DNA handles with biotinylated distal ends.

**Figure S2.**
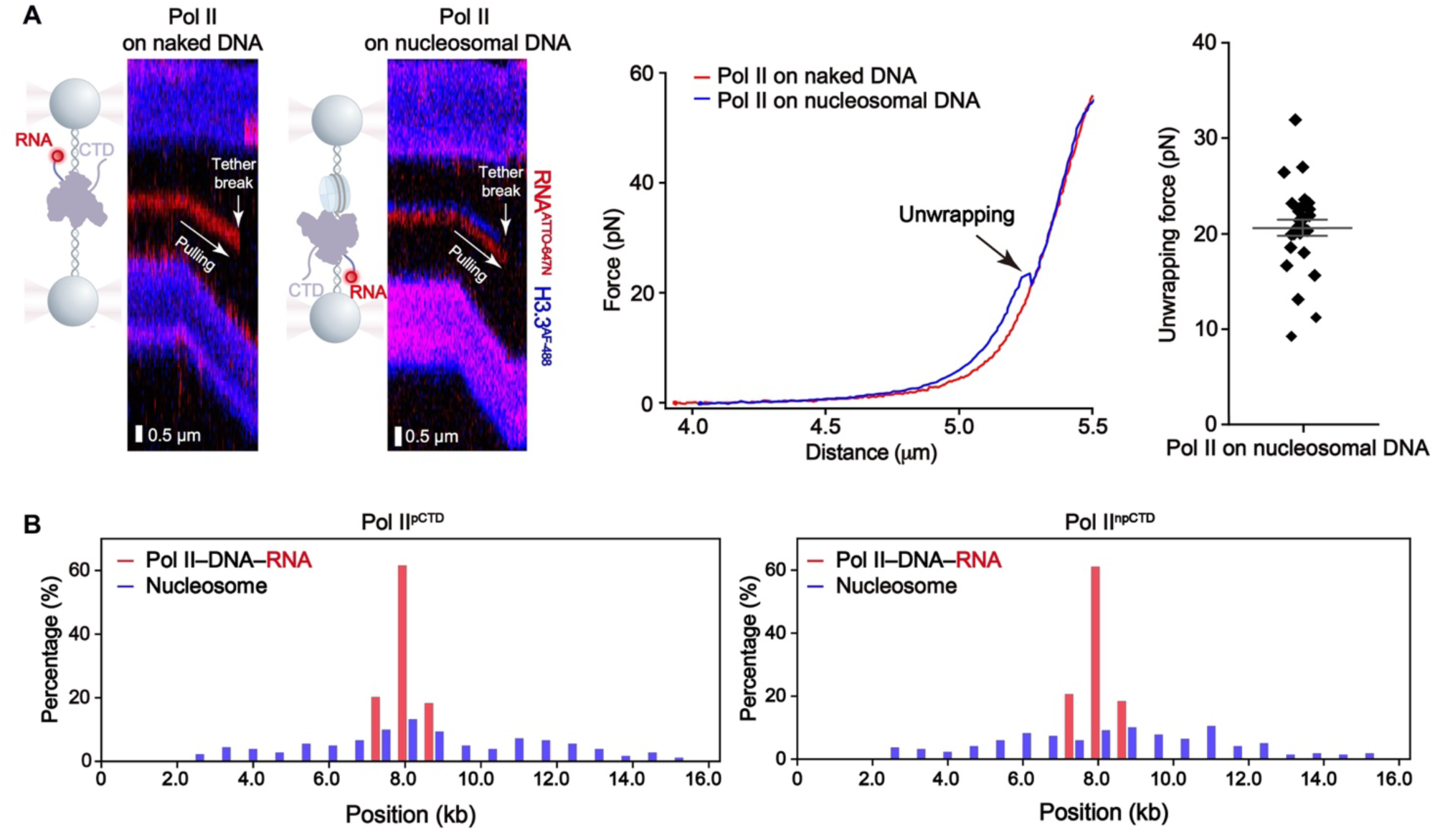
Mechanical validation and spatial distribution of nucleosomes on Pol II-containing DNA tethers. **A**, Comparison of DNA tethers without nucleosomes (naked DNA) and with NAP1-assembled nucleosomes (nucleosome-loaded DNA) [52], both containing preassembled Pol II–DNA–RNA complexes. Left, representative kymographs and schematics (RNA, red; H3.3, blue; scale bars, 0.5 µm). Middle, force–distance curves for naked DNA (red) and nucleosome-loaded DNA (blue). Right, DNA-unwrapping forces (28 measurements from 16 tethers); bars, mean ± s.e.m. **B,** Spatial distributions of preassembled complexes (RNA, red) and nucleosomes (H3.3, blue) before transcription on 16.2-kb tethers containing pre-phosphorylated Pol II (104 tethers) or initially non-phosphorylated Pol II (136 tethers). Positions are measured along the tether; the preloaded complexes lie near its midpoint (approximately 8.1 kb).

**Figure S3.**
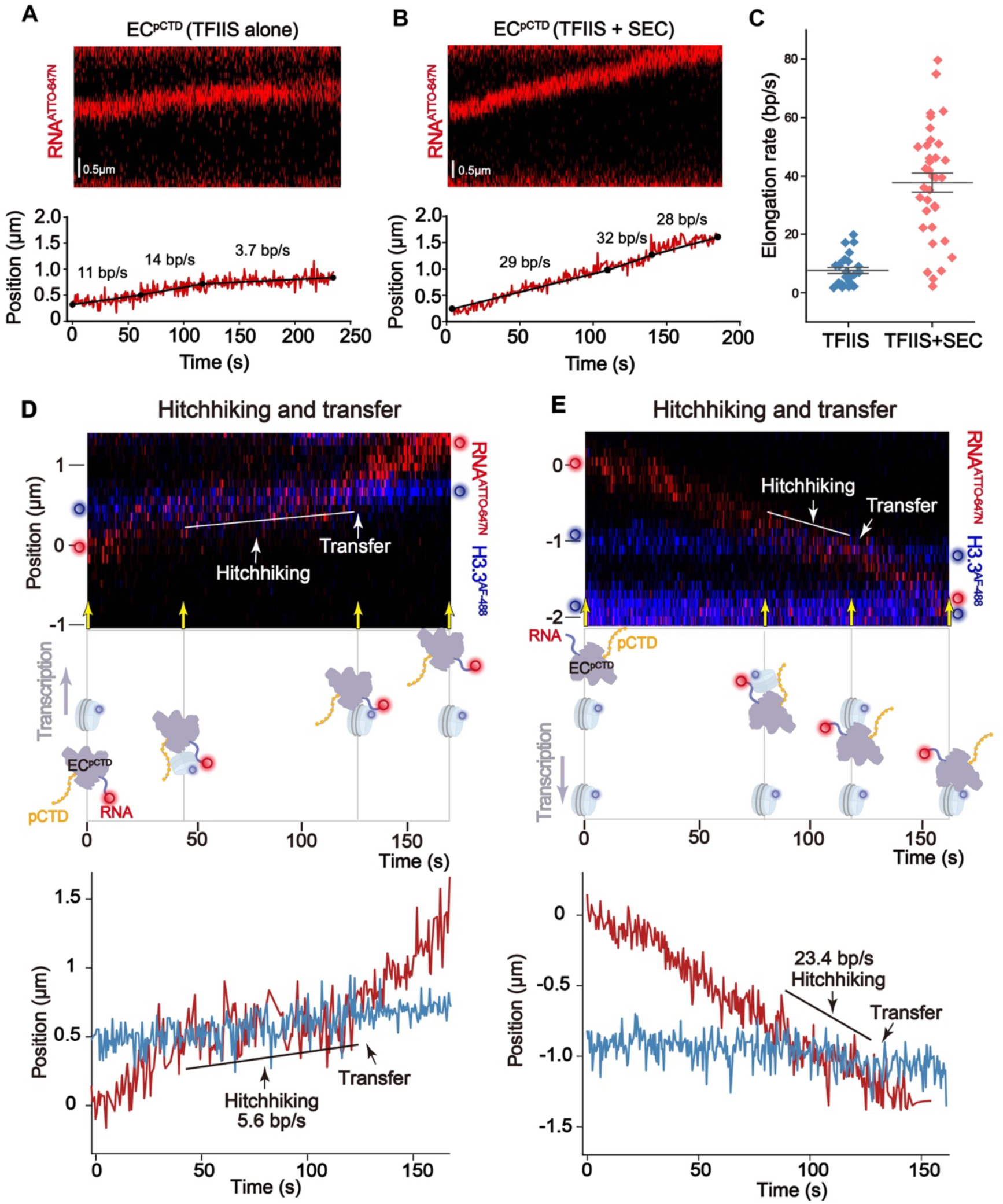
SEC-supported elongation and histone hitchhiking followed by upstream transfer. **A, B,** Representative kymographs (top) and position–time trajectories (bottom) of EC^pCTD^ on naked DNA with TFIIS alone (A) or TFIIS and SEC (B). RNA, red; black lines, piecewise linear fits; filled circles, segment boundaries. Scale bars, 0.5 µm. **C,** Segmental elongation rates with TFIIS alone (26 segments from 10 ECs) or TFIIS and SEC (35 segments from 17 ECs). Each point represents one segment; bars, mean ± s.e.m. **D, E,** Representative EC^pCTD^ trajectories showing slow (D; 5.6 bp/s) and faster (E; 23.4 bp/s) histone hitchhiking followed by transfer to upstream DNA. Top, kymographs; middle, schematic interpretations at the time points indicated by yellow arrows; bottom, corresponding position–time trajectories of ATTO647N-labelled RNA (red) and AF488-labelled H3.3 (blue). White lines in the kymographs and black guide lines in the position–time plots indicate carriage intervals.

**Figure S4.**
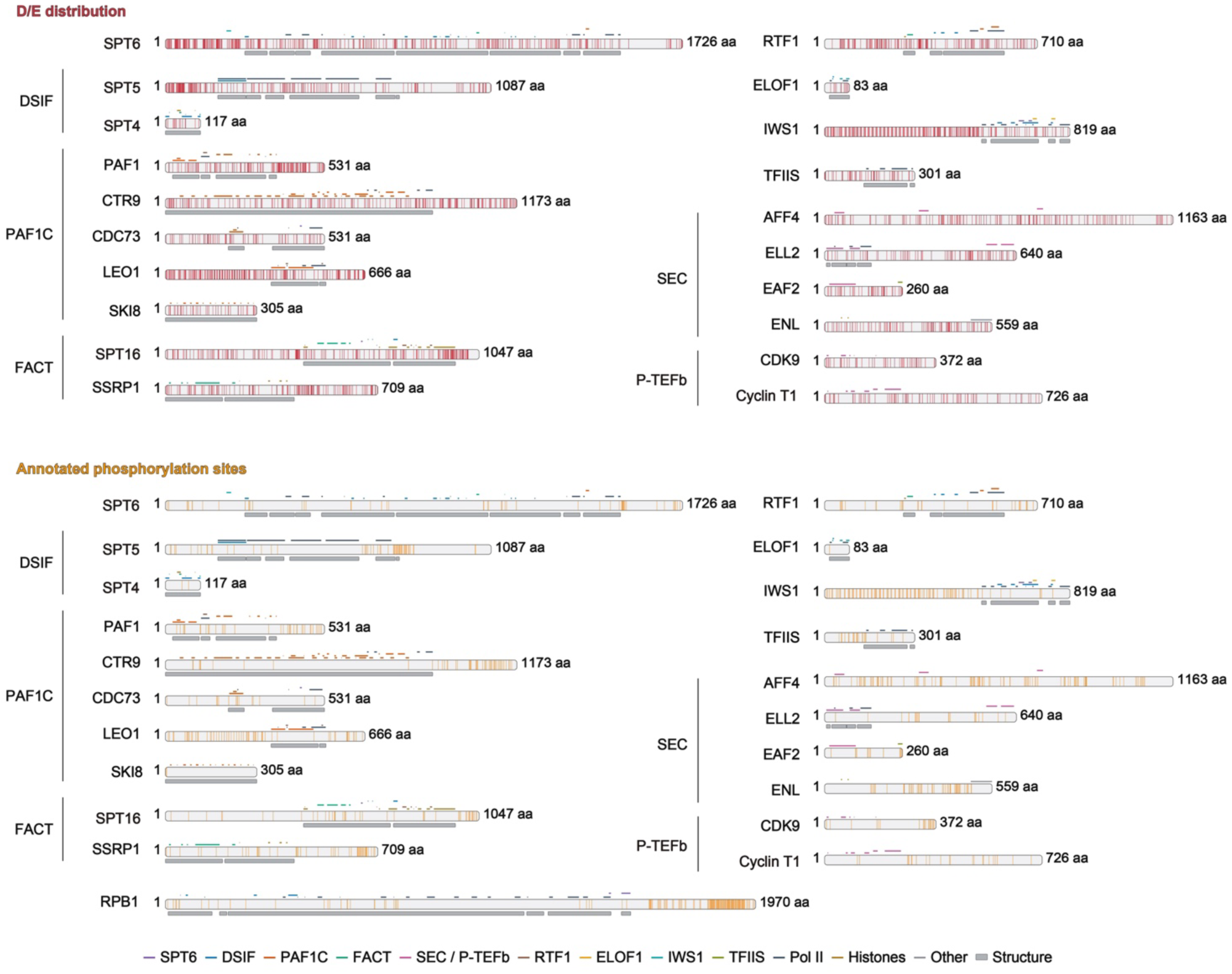
Distribution of acidic residues and annotated phosphorylation sites in human transcription-associated proteins. Sequence maps of acidic residues (aspartate/glutamate; red, upper) and database-annotated phosphorylation sites (orange, lower) in the indicated human transcription-associated proteins. Protein lengths are given in amino acids (aa). Colored lines identify interaction regions and partner proteins or complexes; grey bars indicate residues modelled in selected elongation complex structures. Phosphorylation annotations are from UniProt and iPTMnet; structural and interaction annotations are from PDB structures and published studies. RPB1 is shown in the phosphorylation-site map only.

**Figure S5.**
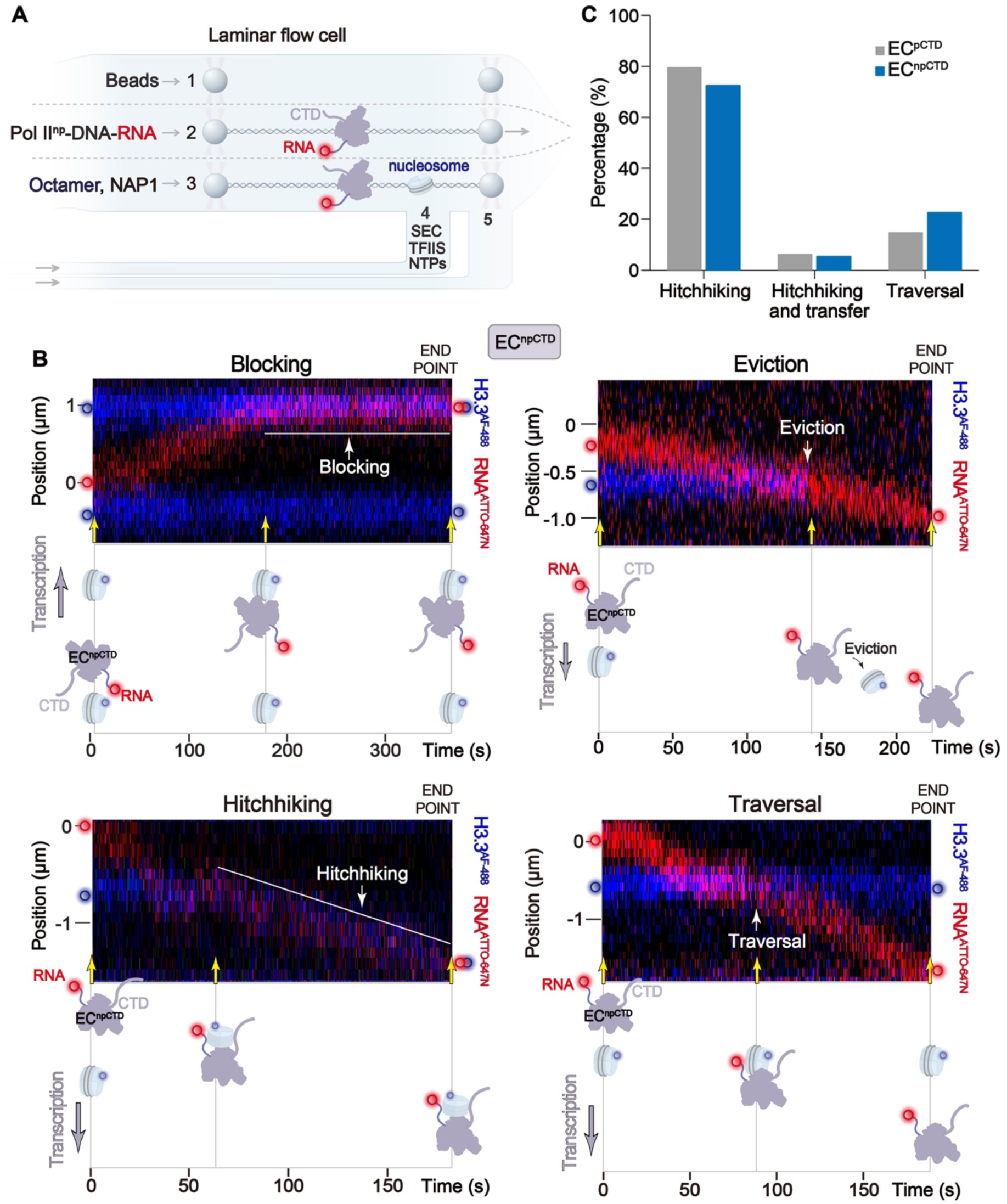
Nucleosome encounters by non-phosphorylated Pol II. **A,** Workflow for nucleosome encounters by EC^npCTD^ with TFIIS and SEC. **B,** Representative blocking, eviction, hitchhiking and traversal records, arranged as in Fig. 1B. **C,** Distributions of continued hitchhiking, hitchhiking and transfer, and traversal in EC^pCTD^ (n = 117 records) and EC^npCTD^ (n = 58 records) after excluding blocking and eviction; percentages are normalized to these three categories.

**Figure S6.**
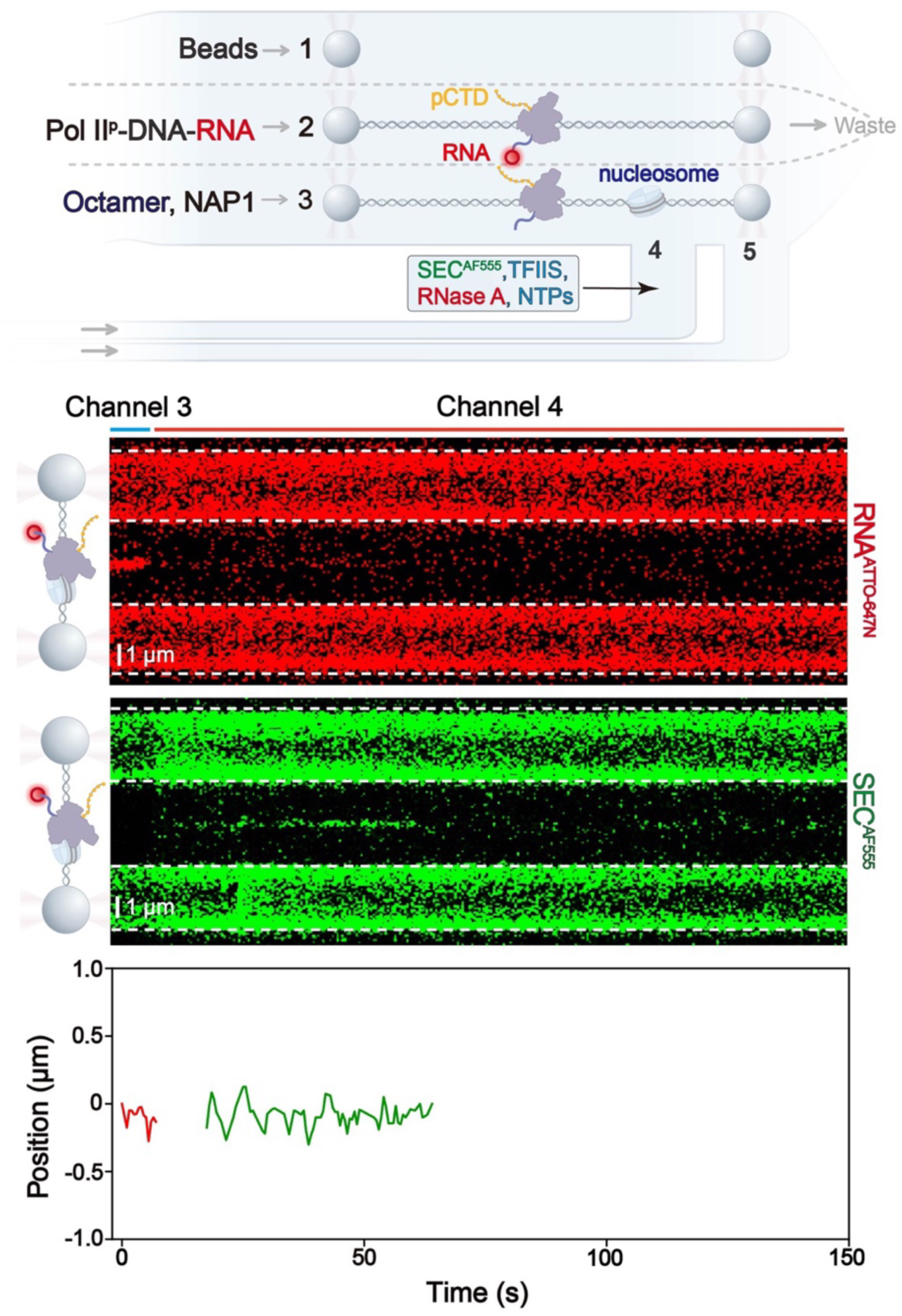
RNase A challenge before nucleosome encounter. Workflow (top), representative kymographs (middle) and extracted trajectories (bottom) for RNase A treatment before nucleosome encounter. Channel 4 contained TFIIS, AF555-labelled SEC, RNase A and NTPs. ATTO647N-labelled RNA, red; AF555-labelled SEC, green. Blue and red bars indicate residence in channels 3 and 4, respectively. Dashed white lines mark bead boundaries. Scale bars, 1 µm. After RNase A treatment, the RNA signal disappears and SEC shows intermittent association near the initial EC position without sustained forward movement.

**Figure S7.**
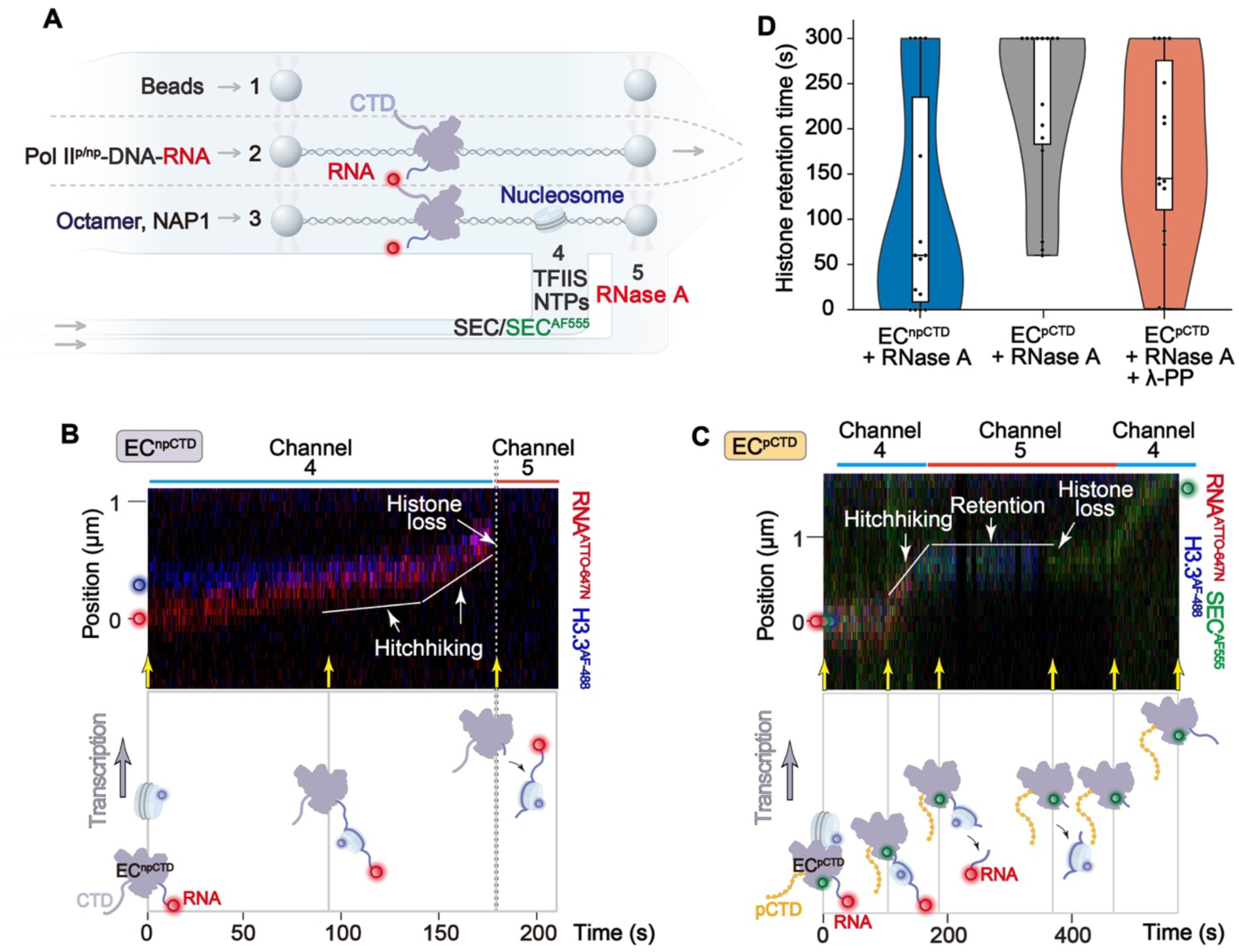
Additional RNase A challenge records and histone-retention times. **A,** Workflow for RNase A challenge after histone carriage. **B,** EC^npCTD^ record showing nearly coincident loss of RNA and histone fluorescence. **C,** EC^pCTD^ record showing histone loss while fluorescent SEC remains and resumes movement on return to the elongation channel. **D,** Observed histone-retention times for RNase A-treated EC^npCTD^, RNase A-treated EC^pCTD^ and RNase A–λ-PP-treated EC^pCTD^, from left to right (n = 15 each; same records as Fig. 2E). Dots, individual records; white boxes, interquartile ranges; central lines, medians; whiskers, minimum and maximum recorded values. Values of 300 s are right-censored; 0 s denotes loss within the time resolution.

**Figure S8.**
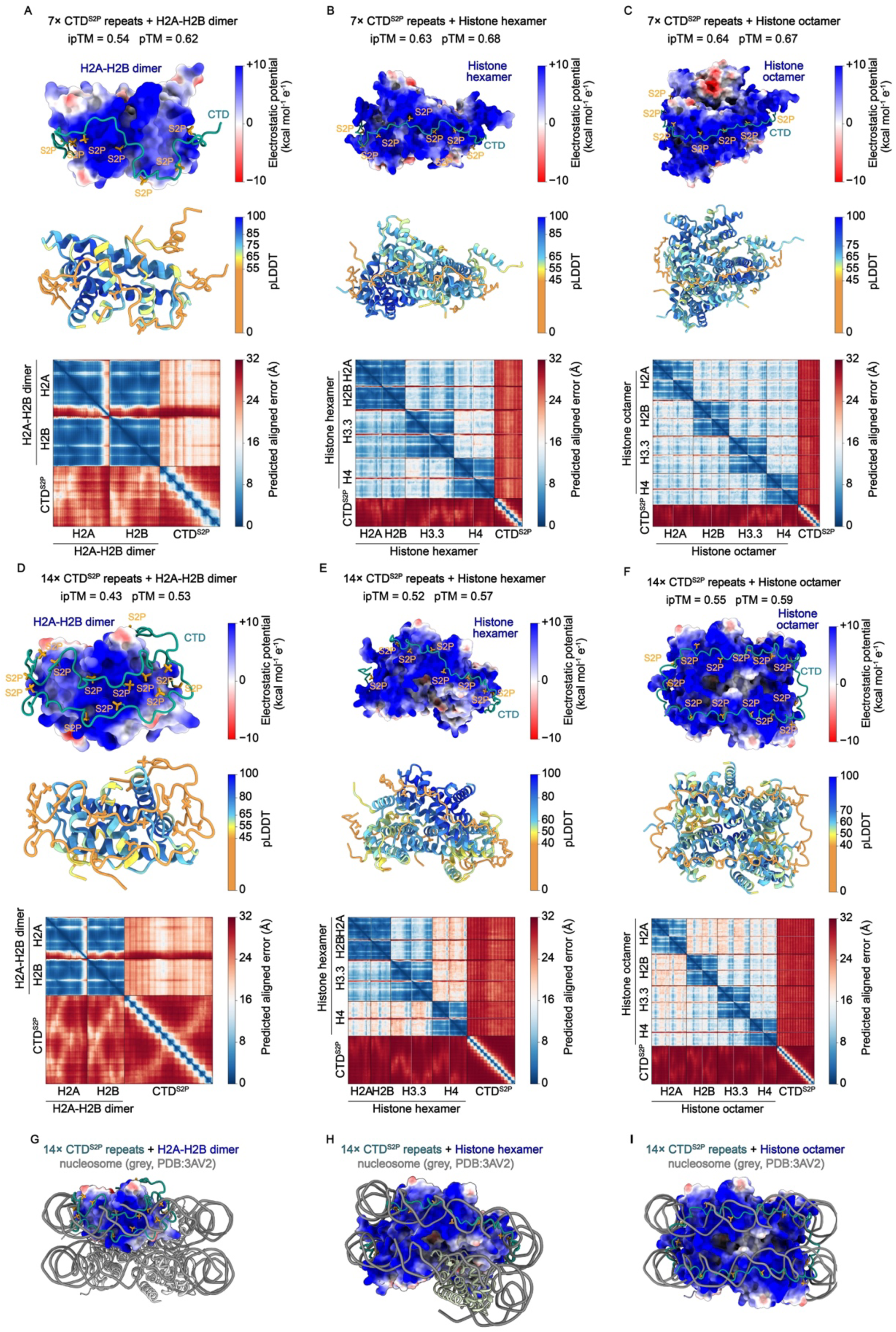
Predicted structures of Ser2-phosphorylated CTD–histone complexes. **A–F,** AlphaFold 3 models of seven (A–C) or fourteen (D–F) Ser2-phosphorylated CTD repeats with an H2A–H2B dimer (A,D), histone hexamer (B,E) or histone octamer (C,F), selected from 25 predictions per condition. Top, histone electrostatic potential (−10 to +10 kcal mol−1 e−1); CTD, teal; Ser2 phosphates, gold. Middle, predicted local distance difference test (pLDDT) scores, colored according to the scale shown in each panel. Bottom, predicted aligned error (PAE; Å). Global interface predicted template modelling (ipTM) and predicted template modelling (pTM) scores are indicated. **G–I,** Superpositions of the fourteen-repeat models with an H3.3 nucleosome (grey; PDB 3AV2). DNA was excluded from all prediction inputs. Panels E and H show alternative views of Fig. 2G.

